# High false sign rates in transcriptome-wide association studies

**DOI:** 10.64898/2025.12.19.695550

**Authors:** Peter A. Gerlach, Nikhil Milind, Jeffrey P. Spence, Jonathan K. Pritchard

## Abstract

Transcriptome-wide association studies (TWAS) are widely used to identify genes involved in complex traits and to infer the direction of gene effects on traits. However, despite their popularity, it remains unclear how accurately TWAS recover the true direction of a gene’s effect on a trait. Here, we estimate the false sign rate (FSR) of TWAS for plasma proteins, leveraging the expectation that increased gene expression should generally increase protein expression. We then extend this framework to complex traits, where loss-of-function burden tests provide the expected direction-of-effect. In both analyses, we observe high discordance with expectations, with TWAS showing an FSR of 23% for plasma proteins and 33% for complex traits. While colocalization-based filtering reduced the FSR, substantial discordance remained, and with substantial loss of recall. However, when we restricted gene-direction assignments for plasma proteins to using only relevant tissues in combination with colocalization-based filtering, the FSR dropped to 11%, and to just 5% if we excluded brain-specific proteins. We propose that much of the sign discordance arises when eQTLs in non–trait-relevant tissues tag GWAS-associated haplotypes via distinct, tightly-linked regulatory variants, yielding spurious TWAS associations with the correct genes but with unreliable direction-of-effect. These findings show that TWAS-based direction-of-effect estimates should be interpreted with caution and raise concerns about the reliability of TWAS more broadly.

## Introduction

Genome-wide association studies (GWASs) have uncovered thousands of genetic associations with complex traits, but it remains challenging to connect these trait-associated variants with their underlying causal mechanisms. Transcriptome-wide association study (TWAS) approaches, such as FUSION [1], PrediXcan [2], and related methods such as SMR [3], are widely used to identify genes that may be driving the GWAS signal at significant loci (Figure 1A) [4–17]. By aggregating variant effects into gene-level signals, these methods provide more mechanistically interpretable information about the gene’s effect on a trait.

**Figure 1.**
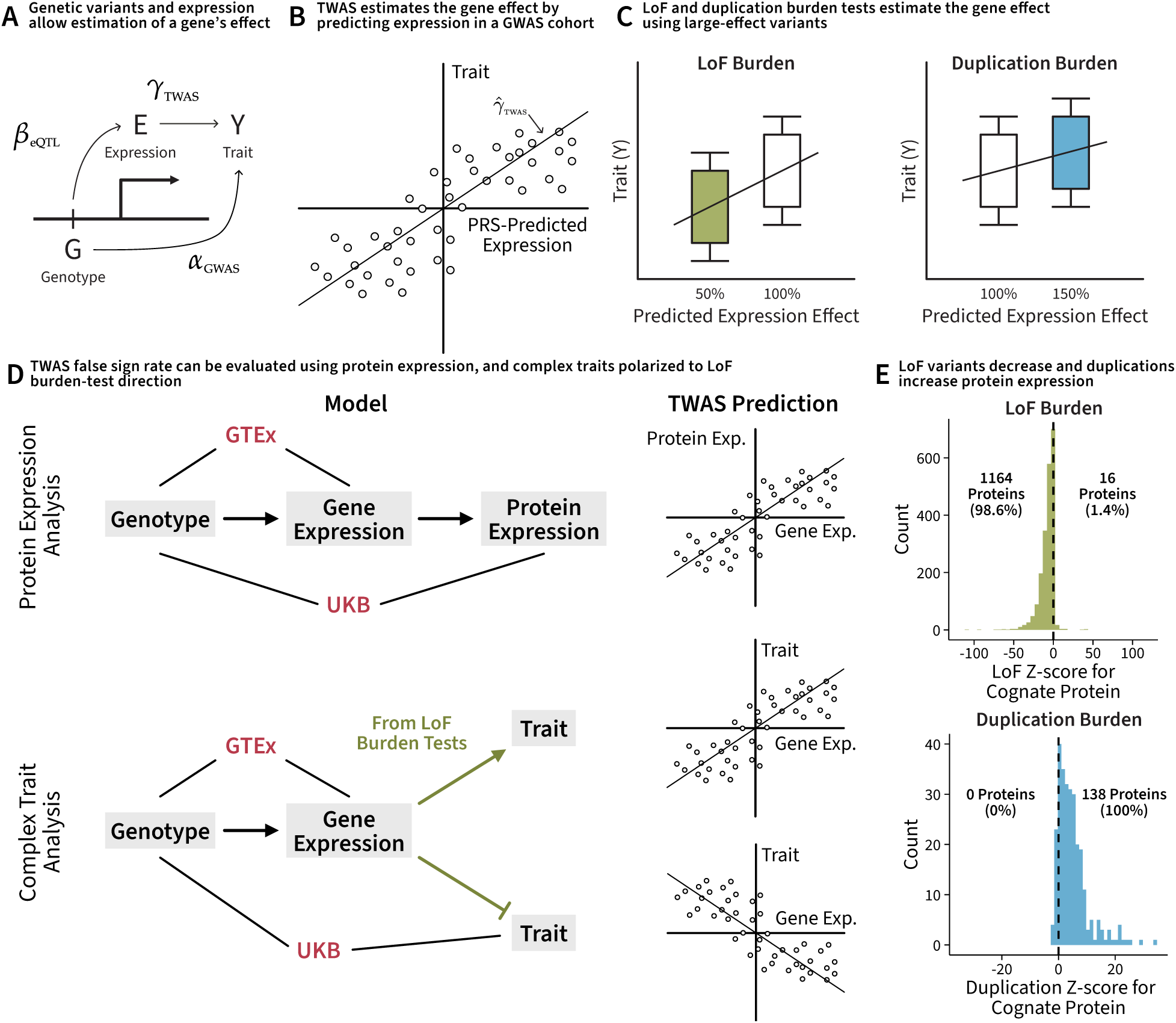
Workflow to evaluate the false sign rate of TWAS. **A.** TWAS estimates the sign and magnitude of effect of a gene on a trait. **B.** TWAS works by using SNPs in the cis region to build a polygenic risk score (PRS) for gene expression, and deploying it in a genome-wide association study (GWAS) cohort to estimate the correlation between gene expression and trait value. **C.** Rare variants provide a gold-standard method to estimate the effect of a gene. LoF variants reduce gene expression, while duplications increase gene expression. We can measure their effect on a trait using burden tests. **D.** We used PRS models for gene expression trained in GTEx, and deployed them in the UKB. We expect protein levels to have positive correlations with gene expression. For complex traits, we used the LoF burden test to determine which direction we expected from TWAS. We used these expectations to compute the FSR. **E.** For plasma proteins in the UK Biobank, LoF variants decrease protein levels, and duplications increase protein levels. We report the number of LoF and duplication burden tests with significant effects on their cognate protein (Bonferroni-corrected p-value less than 0.05) on either side of the dashed line. This makes plasma protein levels an attractive phenotype to assess the false sign rate (FSR).

Conceptually, TWAS leverages genetic variants associated with gene expression (eQTLs) to infer each individual’s genetically predicted expression level for a gene, and then tests whether variation in the genetically predicted expression is associated with variation in the trait. In practice, this involves first building an expression prediction model using a dataset with both genotypes and expression data (*e.g.*, GTEx [18]), and then deploying this model in a dataset where genotypes and traits are available (*e.g.*, UK Biobank [19]). By estimating the association between genetically predicted expression and the trait of interest, TWAS estimates both the magnitude and direction-of-effect of gene expression on the trait (Figure 1B). Estimating the direction of gene effects distinguishes TWAS from other methods that aggregate GWAS signal at the gene level [20, 21] and is critical for downstream tasks, such as inferring coherent gene regulatory networks or interpreting the mechanisms of trait-associated genes for use in drug target discovery [22].

Multiple independent evaluations of TWAS have been conducted, but all of them have focused on the false discovery rate for causal genes [23–27]. Here, we instead focus on characterizing how well TWAS recovers the sign of the gene’s effect on a trait. As we will show, this allows us to detect many instances of spurious associations with the correct genes.

To evaluate the direction-of-effect estimates from TWAS, we assessed the false sign rate (FSR), which is the rate of disagreement between the TWAS-estimated effect direction and the biologically-expected direction. Plasma proteomics provides a unique opportunity to evaluate TWAS direction-of-effect accuracy, because gene expression is expected to increase the abundance of its encoded plasma protein (Figure 1D). Furthermore, we evaluated TWAS performance on predicting the direction-of-effect for various complex traits. Here, we used the loss-of-function (LoF) effect di-rection as the ground truth to determine our expected gene effect direction for TWAS (Figure 1D).

Our results show that even in the favorable setting of plasma proteins, TWAS frequently assigns the wrong direction, and that discordance is even more pronounced for complex traits benchmarked against LoF burden tests. We also show that colocalization-based filtering and focusing on trait-relevant tissues can partially reduce the FSR, but many loci still show confidently incorrect directions that are not well-explained by simple genomic or gene-level features. Together, these findings demonstrate that current TWAS implementations can be highly unreliable for directional inference, and raises questions about whether TWAS and colocalization methods behave as they are commonly understood.

## Results

### Discordant TWAS direction-of-effect estimates for plasma proteins

To evaluate the FSR of TWAS direction-of-effect estimates, we used plasma protein expression as a positive control, since higher gene expression is generally expected to increase protein expression. Specifically, we expect all confident TWAS predictions of plasma protein to be positive (Figure 1D). To test this expectation, we used LoF and duplication burden tests, which are considered a “gold standard” approach to determine the expected gene effect direction (Figure 1C). We indeed observed that 98.6% of proteins that are significant in LoF burden tests show decreased protein expression, and 100% of proteins that are significant in duplication burden tests show increased protein expression (Figure 1E). Together, these results support the expectation that gene expression is generally positively associated with protein expression.

To quantify the FSR of TWAS direction-of-effect estimates for plasma proteins, we applied the FUSION TWAS framework [1] for 2,923 plasma proteins from the UK Biobank Pharma Proteomics Project (UKB-PPP) [28]. We excluded 16 proteins due to either poor quality or because multiple genes encoded the protein, resulting in 2,907 plasma proteins for further analysis (Methods). For each plasma protein, we estimated the direction-of-effect across 47 tissues based on a Bonferroni-corrected significance threshold, with each tissue assigned to one of 27 tissue categories (Table A.1). Of the 2,907 proteins, 383 (13%) lacked a FUSION SNP-weight prediction model in any tissue and 981 (34%) had no significant associations after Bonferroni correction, leaving 1,543 proteins for FSR estimation (Figure 2A). First, we assigned a direction-of-effect for each gene–tissue category by taking a majority vote of significant associations across tissues within that category, and then assigned a gene-level direction by taking a majority vote across tissue categories. We then calculated the FSR across all genes that had at least one significant TWAS association in any tissue. Notably, the estimated FSR was 23%, implying that for roughly one in four genes higher genetically predicted expression is associated with lower plasma protein levels, contrary to the expectation that increased gene expression typically increases protein abundance.

**Figure 2.**
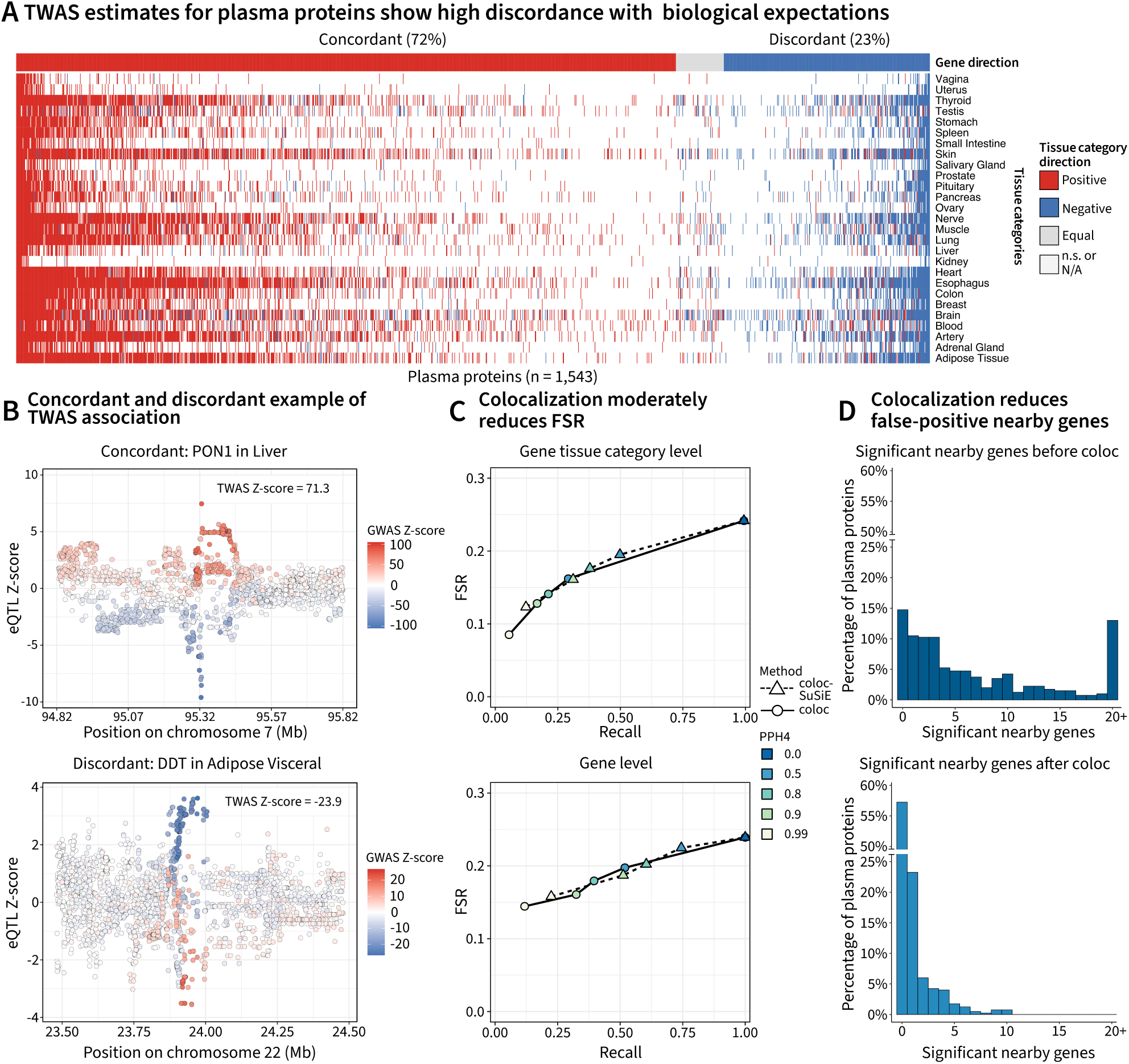
High discordance for TWAS direction-of-effect estimates for plasma proteins. **A**, Heatmap of TWAS direction-of-effect estimates for 1,543 plasma proteins (columns) across 27 tissue categories (rows). For each gene × tissue category, we assigned a direction by majority vote of significant (Bonferroni-corrected) Z-scores across tissues within that category. Tissue categories with a majority of positive Z-scores are colored red, majority negative blue, equal numbers of significant positive and negative grey, and categories with no significant Z-scores or no expression prediction model are white. Gene-level directions were assigned by majority vote across tissue categories, and genes are ordered by concordant (left) versus discordant (right) directions. **B–C**, Example loci illustrating concordant and discordant associations. Points show eQTL Z-scores (y-axis) versus genomic position, with variants colored by their GWAS Z-score. **D–E**, False sign rate (FSR) versus recall for (**D**) gene × tissue-category directions or (**E**) gene-level directions after filtering by PPH4 using coloc (circles, solid line) or coloc-SuSiE (triangles, dashed line). **F–G**, Distribution of the number of TWAS-significant genes within ±2 Mb of each protein-encoding gene (**F**) before and (**G**) after filtering on PPH4 *≥* 0.5, among proteins with at least one colocalizing TWAS association.

To illustrate these patterns, we highlight examples of one concordant and one discordant gene-tissue association (Figure 2B). In the concordant case of *PON1*, we observe that variants that increase expression of *PON1* in the liver also increase plasma protein levels of paraoxonase 1 (PON1) in the blood, as would be expected. In contrast, for the discordant case of *DDT*, we observe that variants that increase *DDT* expression are associated with lower plasma D-dopachrome tautomerase (DDT) levels.

Previous studies have proposed using colocalization-based filtering of TWAS associations to reduce false positives [29], prioritizing genes whose eQTL and GWAS signals are consistent with a shared causal variant. To test whether adding a colocalization filter improved the FSR, we used coloc [30] to calculate the posterior probability that the eQTL and GWAS associations are driven by a shared casual variant, known in coloc as the posterior probability of hypothesis 4 (PPH4). Furthermore, because the standard coloc model assumes a single causal variant per locus, we additionally used coloc-SuSiE, which couples coloc with SuSiE fine-mapping to allow multiple causal variants [31]. coloc supports testing each independent fine-mapped signal one-at-a-time [31], allowing us to potentially increase the recall of TWAS associations. Through TWAS simulations, we show that coloc can indeed filter out false positives and coloc-SuSiE can increase recall (Appendix B).

We calculated recall and FSR for different thresholds of PPH4 from coloc and coloc-SuSiE at the gene–tissue-category level and at the gene level (Figure 2C). Recall was defined as the proportion of significant TWAS associations retained after coloc/coloc-SuSiE PPH4 filtering, relative to the unfiltered set of significant associations with available PPH4 estimates (excluding associations without PPH4). We found that filtering on PPH4 improved the FSR at both the gene–tissue-category and gene levels, with stricter thresholds yielding lower FSR (Figure 2C). However, even at the strictest threshold (PPH4 = 0.99), the gene-level FSR remained above 0.1. Moreover, filtering on PPH4 led to a substantial drop in recall, and although coloc-SuSiE improved recall relative to coloc, it also yielded higher FSR at matched PPH4 thresholds. Thus, colocalization-based filtering only partially resolves discordant TWAS directions for plasma proteins, and a substantial fraction of associations still appear to have incorrect signs even under stringent PPH4 thresholds.

TWAS has been shown to often identify multiple nearby genes at a locus, many of which are likely non-causal “tagging” genes whose predicted expression is correlated with that of the true causal gene due to linkage disequilibrium (LD) or co-regulation [23, 32]. Because plasma levels of a given protein are primarily determined by its encoding gene, expression variation in nearby genes is generally not expected to directly influence that protein’s abundance. This setting therefore provides a convenient ground truth to quantify how many nearby TWAS-significant genes are likely false positive tagging genes. To evaluate how colocalization filtering affects nearby signals, we compared the number of TWAS-significant genes within ±2 Mb of each protein-encoding gene before and after applying coloc filtering at PPH4 *≥* 0.5 (Figure 2D). We observe that prior to colocalization filtering, 85.3% of protein-encoding genes showed at least one TWAS-significant nearby gene, suggesting that non-causal tagging signals are highly prevalent (Figure 2D; top). After colocalization filtering we observed a substantial decrease in nearby significant genes (Figure 2D; bottom), with 43.3% of protein-encoding genes showing at least one TWAS-significant nearby gene. These results indicate that colocalization filtering can markedly reduce nearby TWAS signals that are likely to be non-causal, although many nearby associations persist even after filtering.

### Complex traits show higher discordance of TWAS direction-of-effect estimates compared to plasma proteins

After assessing the FSR on plasma protein levels, we performed the same evaluation on complex traits by identifying 85 quantitative traits (Table A.2) for which GWAS summary statistics from the UKB were publicly available. For these traits, we curated “gold standard” gene-trait pairs from exome-wide significant LoF burden tests, using the sign from the burden test to assign the true sign label to each gene-trait pair. We ascertained 730 gene-trait pairs using the LoF burden tests, which we evaluated using TWAS. While a few genes might have non-monotone gene dosage response curves (*i.e.*, both higher and lower expression of the gene have the same effect on the trait), we expect such curves to dilute the TWAS signal rather than confidently assign the incorrect direction [33].

For 484 of these gene-trait pairs, TWAS identified at least one significant association in a tissue (Figure 3A). Since it is unclear which tissue is causal for any given complex trait or gene, we used the same majority vote strategy as the plasma proteins to summarize the sign for each tissue category and gene. We found that, for a substantial fraction of gene-trait pairs (33%), the gene effect across tissue categories was in the wrong direction. For some gene-trait pairs at the far right of the heatmap, TWAS was strikingly confident in the incorrect association direction in multiple tissues.

**Figure 3.**
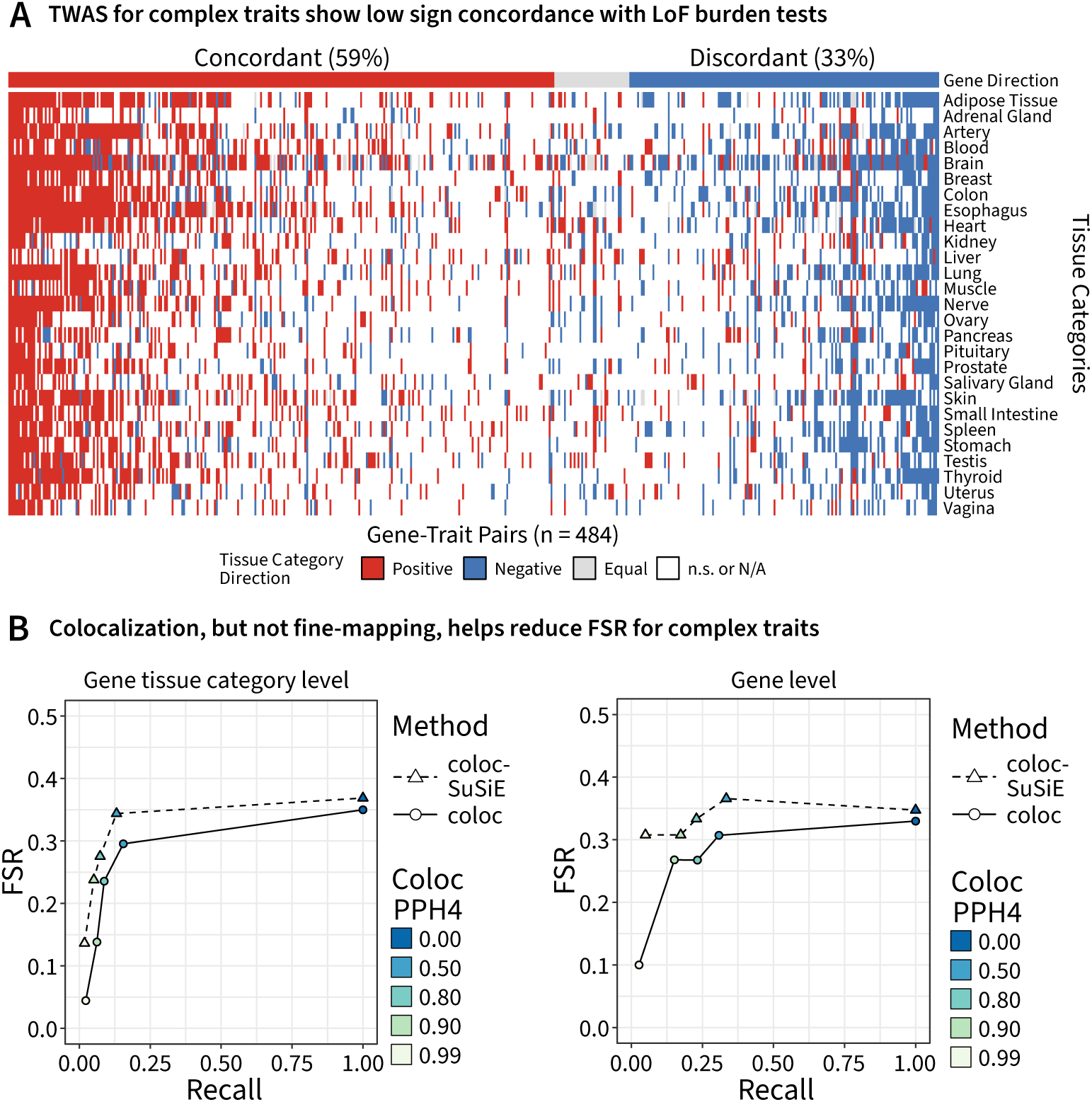
TWAS has a high FSR for complex traits. **A.** Heatmap of TWAS direction-of-effect estimates for 484 gene-trait pairs (columns) across 27 tissue categories (rows). For each gene-trait pair and tissue category, we assigned a direction by majority vote of significant (Bonferroni-corrected) Z-scores across tissues within that category. Tissue categories with a majority of positive Z-scores are colored red, majority negative blue, equal numbers of significant positive and negative grey, and categories with no significant Z-scores or no expression model are white. Gene-level directions were assigned by majority vote across tissue categories, and genes are ordered by concordant (left) versus discordant (right) directions. **B-C.** False sign rate (FSR) versus recall for gene-trait pairs and tissue category directions (**B**) or gene-level directions (**C**) after filtering by PPH4 using coloc (circles, solid line) or coloc-SuSiE (triangles, dashed line).

We again evaluated the use of colocalization and fine-mapping to improve the FSR (Figure 3B). Similar to the proteins, using colocalization as a filtering strategy did reduce the FSR, although with a substantial loss of recall. Fine-mapping again resulted in a higher FSR, but with no concomitant increase in recall.

We built a simple linear model to explore why some gene-trait pairs had a substantially higher FSR than others (Figure 3A) using gene density, local recombination rate, gene constraint measured by *s*_het_, mean *cis*-heritability of the gene, and estimates of trait monotonicity. We accounted for the fact that some genes appear in multiple gene-trait pairs by using a random intercept for the trait and including the sample size for the trait’s GWAS as a fixed effect to account for power differences. Although the model fit the data well based on a likelihood ratio test against a null model with only a random intercept and the sample size (*p* = 3.6 *×* 10*^−^*^4^), implying that the predictors explained some of the variance in FSR, none of the individual predictors were convincingly associated with FSR (Appendix D). This suggests that there is no single gene-level factor among those that we tested that explains why some loci demonstrate striking discordance between TWAS and LoF burden test results.

### Misleading TWAS signals in non–trait-relevant tissues

Previous work has shown that gene–trait associations are often tissue specific, motivating us to ask whether focusing on a trait-relevant tissue for each plasma protein could improve the direction-of-effect FSR compared to a majority vote across tissue categories [23, 29]. Using tissue-of-origin labels from the Human Proteome Distribution Atlas [34], we restricted to proteins specific to a single tissue, resulting in 228 proteins for which we defined gene direction using the TWAS Z-score in that tissue. Overall, 20% of proteins were discordant using this approach (Figure 4A), compared to 23% when taking a majority vote across tissues (Figure 2A). Notably, we observed particularly high discordance for brain-specific proteins, and excluding these reduced discordance to 14%. Applying coloc filtering at PPH4 *≥* 0.5 further reduced discordance to 11%, and to just 5% after additionally removing brain-specific proteins (Figure 4B). Together, these results suggest that combining trait-relevant tissue information with colocalization-based filtering can substantially improve the directional accuracy of TWAS effect estimates for plasma proteins, although we recognize that identifying the relevant tissues for complex traits remains an open challenge [23, 29, 35].

**Figure 4.**
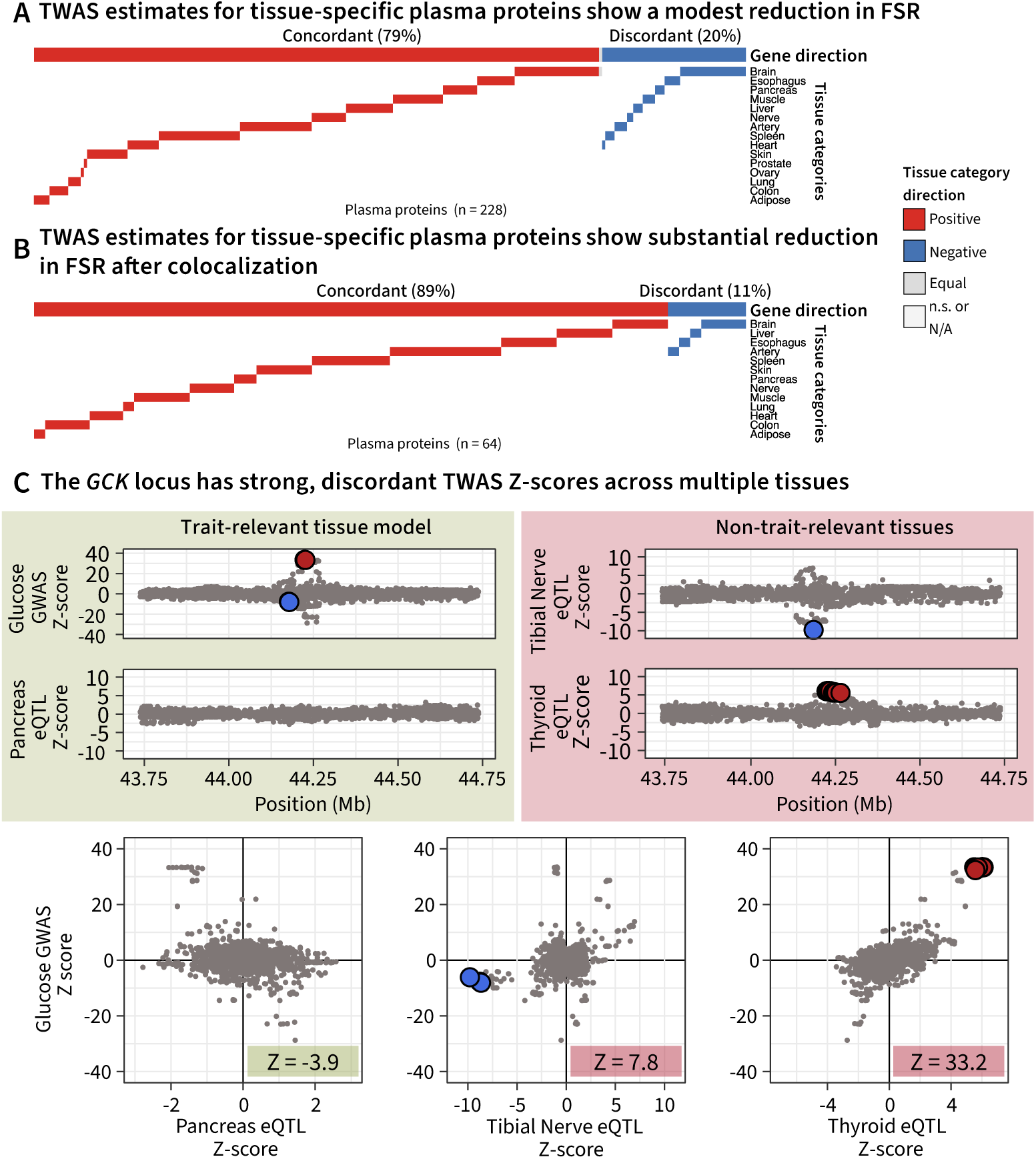
TWAS shows improved direction-of-effect estimates in trait-relevant tissues. A-B, Heatmap of TWAS direction-of-effect estimates for tissue-specific plasma proteins (colums), restricted to their assigned tissues (rows) based on annotations from Malmström et al. [34]. Tissue categories with a majority of positive Z-scores are colored red, majority negative blue, equal numbers of significant positive and negative grey. Genes are ordered by overall sign (positive, neutral, negative), and tissues (rows) are ordered by descending tissue FSR (highest FSR at the top). (**A**) shows TWAS estimates with no colocalization filtering (n = 228 genes) and (**B**) shows TWAS estimates after coloc filtering at PPH4 *≥* 0.5 (n = 64 genes). **C.** *GCK* is an example locus that shows strong discordance across multiple tissues. We expect a negative Z-score in pancreas, but see little evidence of eQTL signal there. In contrast, we see colocalization with eQTL in tibial nerve and thyroid, which are strong but in the wrong direction. Blue and red points are 95% credible sets that colocalize between the eQTl and GWAS data.

To illustrate these tissue-specific discrepancies in complex traits, we examined the association between *GCK* and blood glucose levels as an example. *GCK* encodes glucokinase (GCK), an enzyme that catalyzes the first step in glycolysis and acts as a glucose sensor in various tissues. LoF variants in *GCK* are associated with increased blood glucose levels [36–38]. The mechanism is thought to occur via pancreatic beta cells, where higher *GCK* increases insulin secretion and lowers blood glucose [39, 40]. Therefore, for this gene, we expect TWAS to have a negative Z-score in pancreas, since increased *GCK* expression should be associated with reduced glucose levels. We found that even though there is a strong GWAS signal present at the *GCK* locus, there was no eQTL signal in the pancreas, resulting in no significant TWAS association in the tissue. However, visually, the direction of association in the pancreas appears to be correct (Figure 4C), and using the top eQTL as the TWAS model results in a nominally-significant association in the correct direction (Z = -3.9).

Surprisingly, *GCK* showed significant positive TWAS associations in 17 other tissue categories, even after coloc filtering at PPH4 *≥* 0.5. Paradoxically, the most prominent signals arose in tissues with no obvious mechanistic link to glucose homeostasis. The strongest *GCK* eQTL signal was in tibial nerve, where variants produced a significant positive TWAS association (Z = 7.8) (Figure 4C). Additionally, we observed striking positive TWAS association between thyroid eQTL and glucose levels (Z = 33.2). We detected a single credible set in both tissues, which colocalized with two separate glucose GWAS credible sets (PPH4 = 0.51 and PPH4 = 0.99 in tibial nerve and thyroid respectively). Taken together, the locus-level patterns in these tissues provide strong evidence that the same variants underlie both the eQTL and GWAS signals, but with an inferred direction-of-effect opposite to that expected from *GCK* biology (Figure 4C). We explored some additional loci with similarly discordant associations across tissue groups, with varying colocalization evidence (Appendix C), many showing similarly strong evidence in the wrong direction.

Previous studies have suggested prioritizing tissues with the largest eQTL sample sizes, arguing that *cis*-regulated gene expression levels exhibit high genetic correlation across tissues and that well-powered proxy tissues can therefore substitute for the causal tissue when it is difficult to assay [32]. In contrast, our results show that relying on non–trait-relevant tissues in this way can yield highly significant but directionally misleading TWAS signals, with loci such as *GCK* displaying strong, colocalizing associations in the wrong direction. Given these results, we hypothesize that these discordant signals in non–trait-relevant tissues arise when GWAS and eQTL signals are driven by distinct variants in very high linkage disequilibrium, which neither TWAS nor colocalization can reliably distinguish. In such settings, genes with little or no causal effect in a non-trait-relevant tissue can nonetheless inherit large TWAS Z-scores, simply because their expression tags the same underlying genetic variation that drives the true causal mechanism in an unobserved or underpowered trait-relevant tissue. We expect such patterns to arise if expression in the trait-relevant tissue is constrained relative to non-trait-relevant tissues, resulting in fewer discoverable eQTL [41]. The eQTL in unconstrained tissues can be in high LD with the relevant eQTL, but have a random sign relative to the trait.

## Discussion

Taken together, we show that TWAS direction-of-effect estimates show high discordance with biological expectations for both plasma proteins (23%) and complex traits (33%). While we found colocalization-based filtering to reduce the FSR, a substantial discordance still remained after filtering. We found that restricting gene-direction assignments to trait-relevant tissues and applying colocalization-based filtering markedly improved the FSR for plasma proteins, reducing discordance to 11%, and to just 5% when excluding brain-specific proteins.

These results are broadly consistent with a failure mode in which the causal eQTL is in high LD with distinct eQTL variants in non–trait-relevant tissues, such that TWAS and colocalization correctly implicate the underlying haplotype but misattribute the tissue and direction-of-effect. We expect these patterns in part because regulatory variants that affect a gene’s expression only in non–trait-relevant tissues are likely to be under weaker purifying selection, and can therefore attain larger effect sizes or higher frequencies than variants acting in trait-relevant tissues [41]. When such relaxed-constraint eQTLs are in high LD with the true causal variant, they can generate significant TWAS signals in non-relevant tissues. By contrast, focusing on trait-relevant tissues restricts attention to the eQTLs that plausibly mediate the biological effect on the trait, thereby testing the relevant eQTLs against the GWAS signal. Moreover, because non–trait-relevant tissues usually far outnumber trait-relevant tissues, these spurious but significant signals could easily end up dominating TWAS results for a gene tested across all tissues.

In addition to this LD-driven failure mode, other biological mechanisms could also contribute to the observed discordance. First, tissue-specific eQTL effects may play a role: while eQTL effect sizes are generally concordant across tissues [42], some variants can exhibit opposite effects in different tissues [43], potentially leading to mismatched gene-level directions when non-relevant tissues are included. Second, non-monotone gene–trait relationships [33] may cause LoF and common-variant effects to differ in sign, though we expect this to be rare. Third, co-regulation of nearby genes can propagate the signal from a truly causal gene to its neighbors, yielding statistically significant but misleading TWAS associations [23, 32]. Finally, a gene may have opposite effects on the trait in different tissues, so that LoF variants, which perturb all tissues at once, produce a net phenotypic effect whose sign can differ from tissue-specific eQTL or TWAS estimates.

Furthermore, technical factors may also partly explain the observed discordance. Because eQTL and GWAS are typically measured in different cohorts, TWAS often relies on an external LD reference panel. At loci where abundant GWAS signal is present, the TWAS model may be confident that an association is present (resulting in a large absolute Z-score), but subtle mismatches in LD patterns may cause sign flipping. In addition, most TWAS frameworks treat predicted expression as fixed, ignoring uncertainty in the expression imputation model, which can induce spurious associations with essentially random sign at strongly associated loci [44].

We recommend that current and future TWAS methods should be evaluated for their FSR in addition to their FDR. Our analysis show that pQTL data sets are an attractive source of phenotypes to test the FSR, since we rarely expect to see a negative relationship between the genetic component of gene expression and protein levels.

While LoF burden tests often act as a “gold standard” for gene sign, ascertainment of burden tests prioritizes genes by their specificity to the trait, suggesting that the genes we considered may not be representative of all genes [45]. Common variant signals from GWAS can be more context specific, which would allow us to infer the sign of genes that may be too constrained to be discovered by burden tests. TWAS methods are invaluable for summarizing the evidence for the sign of a gene’s effect from common variants. Continued development and refinement of these methods will be crucial for understanding the causal, mechanistic role of genes in various traits and diseases.

## Methods

### Burden test summary statistics

We retrieved burden test summary statistics from our prior analysis in the UKB [33]. For the protein burden tests, we retrieved the test results for only the cognate gene.

To identify “top hits” from the LoF burden tests for complex traits, we used a Bonferroni threshold based on the number of genes and number of traits. At the 5% significance level, the corrected p-value threshold was 3.32 *×* 10*^−^*^8^. At this threshold, we identified 730 significant gene-trait pairs in 73 of the 85 chosen complex traits.

### TWAS

For protein expression analyses, we used pQTL summary statistics for participants of European ancestry (n = 34,557) from the UKB-PPP [28]. Of the 2,923 plasma proteins, one protein (GLIPR1) was excluded due to data failing quality control [28], and 15 plasma proteins were excluded due to multiple genes encoding the proteins (CKMT1A_CKMT1B, DEFA1_DEFA1B, DEFB4A_DEFB4B, EBI3_IL27, FUT3_FUT5, IL12A_IL12B, LGALS7_LGALS7B, MICB_MICA, AMY1A_AMY1B_AMY1C, BOLA2_BOLA2B, CGB3_CGB5_CGB8, CTAG1A_CTAG1B, DEFB103A_DEFB103B, DEFB104A_DEFB104B, SPACA5_SPACA5B) resulting in 2,907 plasma proteins used for TWAS associations.

For complex traits, we used summary statistics from the Neale Lab for 85 traits measured in the UK Biobank (Table A.2).

We used FUSION to perform TWAS [1]. We performed TWAS between all genes and traits across all GTEx tissues for which expression models were available. We used the GTEx expression models and 1000 Genomes project linkage disequilibrium (LD) reference panel provided with the FUSION software.

### Annotating gene direction

To determine the overall direction of gene effects, we performed a hierarchical majority vote. First, for each tissue category, we determined the direction by majority vote across its tissues. Next, we performed a majority vote across all tissue categories to assign the final direction for each gene. The grouping of GTEx tissues into categories is provided in Table A.1.

### Colocalization and fine-mapping

We used the susieR package [46, 47] to perform fine-mapping of the eQTL and GWAS summary statistics. We performed fine-mapping on 1 Mbp regions centered at the transcription start site (TSS) for all genes in GENCODE v39 [48]. We used genotypes from individuals assigned to the “EUR” superpopulation in the 1000 Genomes project [49] to construct the LD reference panel. We required that at least 10 variants needed to be present in both the summary statistics and the LD panel to proceed with fine-mapping. We used the susie_rss function to fit the sum of single effects (SuSiE) regression model to the summary statistics and LD panel.

We tested for colocalization using the coloc package [30, 31]. To perform colocalization only, we used the approximate Bayes factor approach implemented in the original package [30], which assumes one causal variant at each locus. To perform colocalization with SuSiE, we tested all pairwise combinations of the independent signals detected between eQTL and GWAS summary statistics using the coloc.bf_bf function. We excluded any loci that did not have at least 100 overlapping variants between the eQTL and GWAS summary statistics from the colocalization analysis.

### Simulation of GWAS summary statistics

We simulated protein expression GWAS summary statistics (pQTLs) in a 1 Mb window around the *MBL2* locus using the sim_mv_determined function from the GWASBrewer package [50, 51]. To approximate the correlation structure observed in UKB-PPP, we used LD matrices and allele frequencies from the UK Biobank computed in Zhao et al. [52].

The sample size was fixed at *N* = 30,000. One or two causal variants (depending on the scenario) were specified, with their per-allele effect parameterized to achieve pre-specified Z-scores, while all other variants were assigned zero direct effect size.

For each of the three scenarios in Figure 3, we generated 50 independent replicates of GWAS summary statistics and performed TWAS, colocalization, and fine-mapping analyses as described above.

### Tissue-specificity

To assess tissue-specificity of TWAS signals (Figure 4E), we focused on genes annotated as tissue-specific to a single tissue in Malmström et al. [34] (Table S13). Further we filtered for Global label score >= 1. For each tissue with more than 20 genes specific to that tissue, we compared the TWAS Z-scores of those genes in that tissue to the TWAS Z-scores of the same genes in all remaining tissues.

## Data Availability

Data generated by this study will be placed on Zenodo (https://doi.org/10.5281/zenodo.17990927). Data processed by this study can be accessed via the following URLs: Protein QTL Summary Statistics (https://www.synapse.org/Synapse:syn51364943/files/); Neale Lab’s GWAS Summary Statistics (http://www.nealelab.is/uk-biobank); Tissue Weights from TWAS Website derived from GTEx v8 (http://gusevlab.org/projects/fusion/); LD from 1000 Genomes Project for FUSION (https://alkesgroup.broadinstitute.org/FUSION/); 1000 Genomes Project VCFs for colocalization and fine-mapping LD Panel (https://ftp.1000genomes.ebi.ac.uk/vol1/ftp/release/20130502/, http://ftp.1000genomes.ebi.ac.uk/vol1/ftp/data_collections/1000_genomes_pro ject/release/20190312_biallelic_SNV_and_INDEL/); GTEx eQTL Summary Statistics for Coloc and SuSiE from the eQTL Catalogue (https://www.ebi.ac.uk/eqtl/); LoF Burden Test Summary Statistics (https://zenodo.org/records/16800547); LD and AF summary from UKB for GWAS simulations (https://uchicago.app.box.com/s/jqocacd2fulskmhoqnasrknbt59x3xkn/folder/234629250877).

## Code Availability

Code will be placed on Zenodo (https://doi.org/10.5281/zenodo.17990927).

## Acknowledgments

We thank H. Mostafavi, T. Gjorgjieva, and H. Zhu for feedback on the manuscript and the members of the Pritchard laboratory for helpful discussions. This work was supported by the National Institutes of Health (R01HG014005, R01HG008140, and U01HG012069). We thank Stanford University and the Stanford Research Computing Center for providing computational resources and support that contributed to this research.

## Author Contributions

## A Appendix 1

**Table A.1.** Tissues from GTEx grouped into tissue categories.

| GTEx tissue | Tissue category |
| --- | --- |
| Adipose Subcutaneous | Adipose Tissue |
| Adipose Visceral Omentum | Adipose Tissue |
| Adrenal Gland | Adrenal Gland |
| Artery Aorta | Artery |
| Artery Coronary | Artery |
| Artery Tibial | Artery |
| Brain Amygdala | Brain |
| Brain Anterior cingulate cortex BA24 | Brain |
| Brain Caudate basal ganglia | Brain |
| Brain Cerebellar Hemisphere | Brain |
| Brain Cerebellum | Brain |
| Brain Cortex | Brain |
| Brain Frontal Cortex BA9 | Brain |
| Brain Hippocampus | Brain |
| Brain Hypothalamus | Brain |
| Brain Nucleus accumbens basal ganglia | Brain |
| Brain Putamen basal ganglia | Brain |
| Brain Spinal cord cervical c-1 | Brain |
| Brain Substantia nigra | Brain |
| Breast Mammary Tissue | Breast |
| Colon Sigmoid | Colon |
| Colon Transverse | Colon |
| Esophagus Gastroesophageal Junction | Esophagus |
| Esophagus Mucosa | Esophagus |
| Esophagus Muscularis | Esophagus |
| Heart Atrial Appendage | Heart |
| Heart Left Ventricle | Heart |
| Kidney Cortex | Kidney |
| Liver | Liver |
| Lung | Lung |
| Minor Salivary Gland | Salivary Gland |
| Muscle Skeletal | Muscle |
| Nerve Tibial | Nerve |
| Ovary | Ovary |
| Pancreas | Pancreas |
| Pituitary | Pituitary |
| Prostate | Prostate |
| Skin Not Sun Exposed Suprapubic | Skin |
| Skin Sun Exposed Lower leg | Skin |
| Small Intestine Terminal Ileum | Small Intestine |
| Spleen | Spleen |
| Stomach | Stomach |
| Testis | Testis |
| Thyroid | Thyroid |
| Uterus | Uterus |
| Vagina | Vagina |
| Whole Blood | Blood |

**Table A.2.**
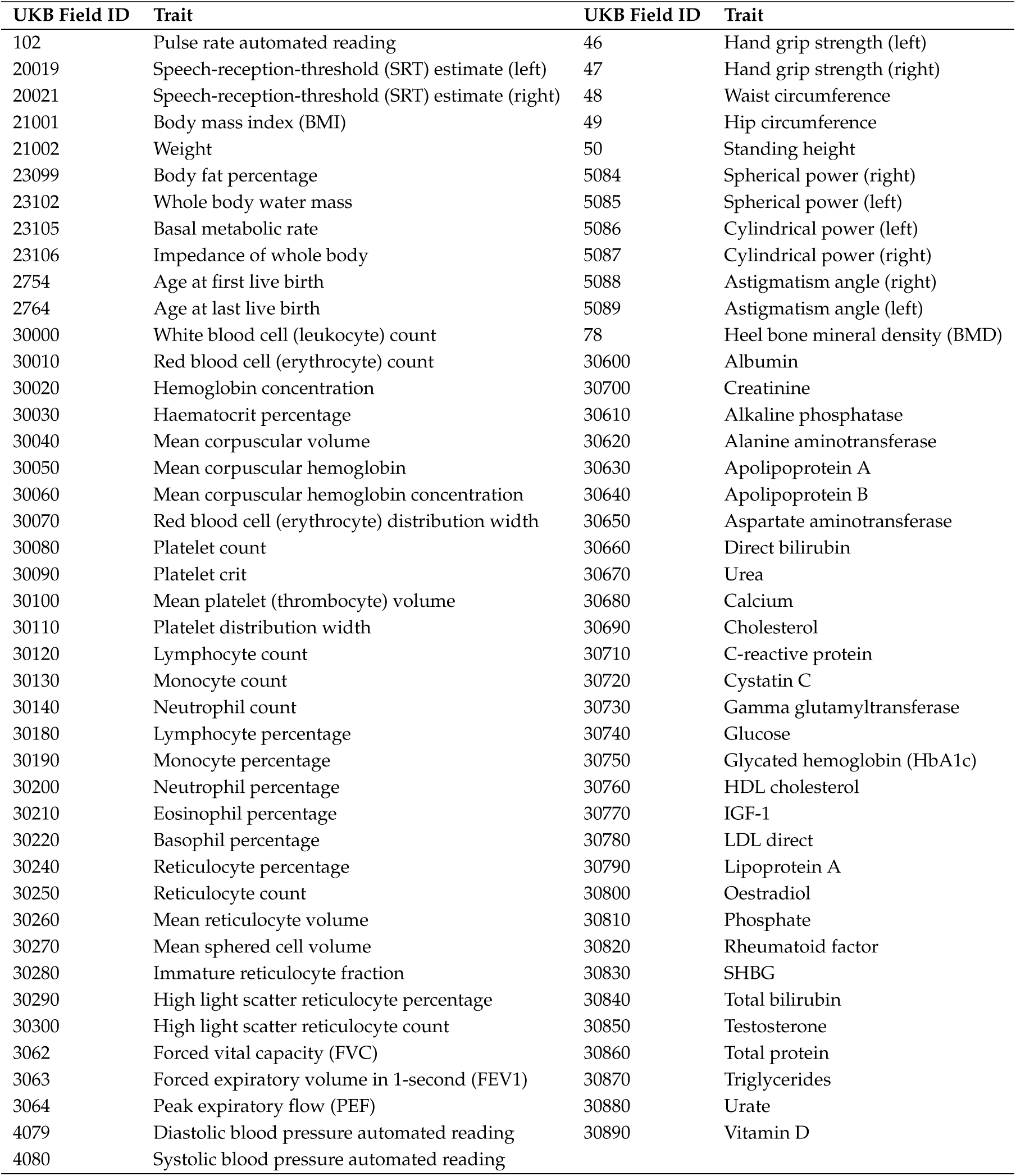
Complex traits used for analysis.

## B Appendix 2

To assess how coloc and coloc.susie affect false positives and recall, we simulated TWAS data under different scenarios. To most realistically simulate the plasma protein TWAS associations from Figure 1, we simulated GWAS summary statistics with the GWASBrewer package [1, 2] using LD matrices and allele frequencies from the UK Biobank computed by Zhao et al. [3] (Figure 3A, 3B, and 3C). Because LD matrices and allele frequencies from GTEx were unavailable, we instead used eQTL summary statistics at the *MBL2* locus in liver tissue, which exhibited a clear signal, and assumed that the top eQTL variant was the causal variant (Figure 3D, 3E, and 3F). We then calculated TWAS Z-scores with FUSION [4] using the provided LD reference from 1000 genomes as similarly done for the plasma proteins (Figure 3A, 3B, and 3C). Finally, we computed PPH4 using both coloc and coloc.susie, and for coloc.susie we report the maximum PPH4 across all SuSiE credible sets (Figure 3G, 3H, and 3I).

In the first scenario, we simulated a “true positive” case in which the causal variant was the same in both eQTL and GWAS summary statistics (Figure 3A, 3D, and 3G). As expected we observe a highly significant TWAS Z-score and a high colocalization probability from both coloc and coloc.susie.

In the second scenario, we simulated the GWAS causal variant to differ from the causal eQTL variant but to be in modest LD with it (*R*^2^ = 0.35) (Figure 3B, 3E, and 3H). Although the TWAS Z-score remained highly significant, both coloc and coloc.susie provided no evidence of colocalization. This indicates that coloc-based methods can correctly filter out “false positive” TWAS signals that arise from LD between distinct causal variants.

In the final scenario, we simulated two distinct GWAS causal variants in which the smaller-effect variant was the same as the eQTL causal variant and the larger-effect GWAS variant being effectively unlinked with the eQTL causal variant (*R*^2^ = 0.002) (Figure 3C, 3F, and 3I). In this scenario we still observe a strong significant TWAS signal. However, because coloc assumes a single causal variant per trait, it attributes the GWAS signal to the larger-effect variant and consequently detects no colocalization with the eQTL causal variant (Figure 3I). Meanwhile, coloc.susie pro-vides strong evidence of colocalization, as it can resolve multiple distinct association signals at the locus and assess colocalization for each pair of signals between traits. Thus, coloc.susie should be able to retain TWAS associations in which smaller effect causal variants colocalize to improve recall.

**Figure B.**
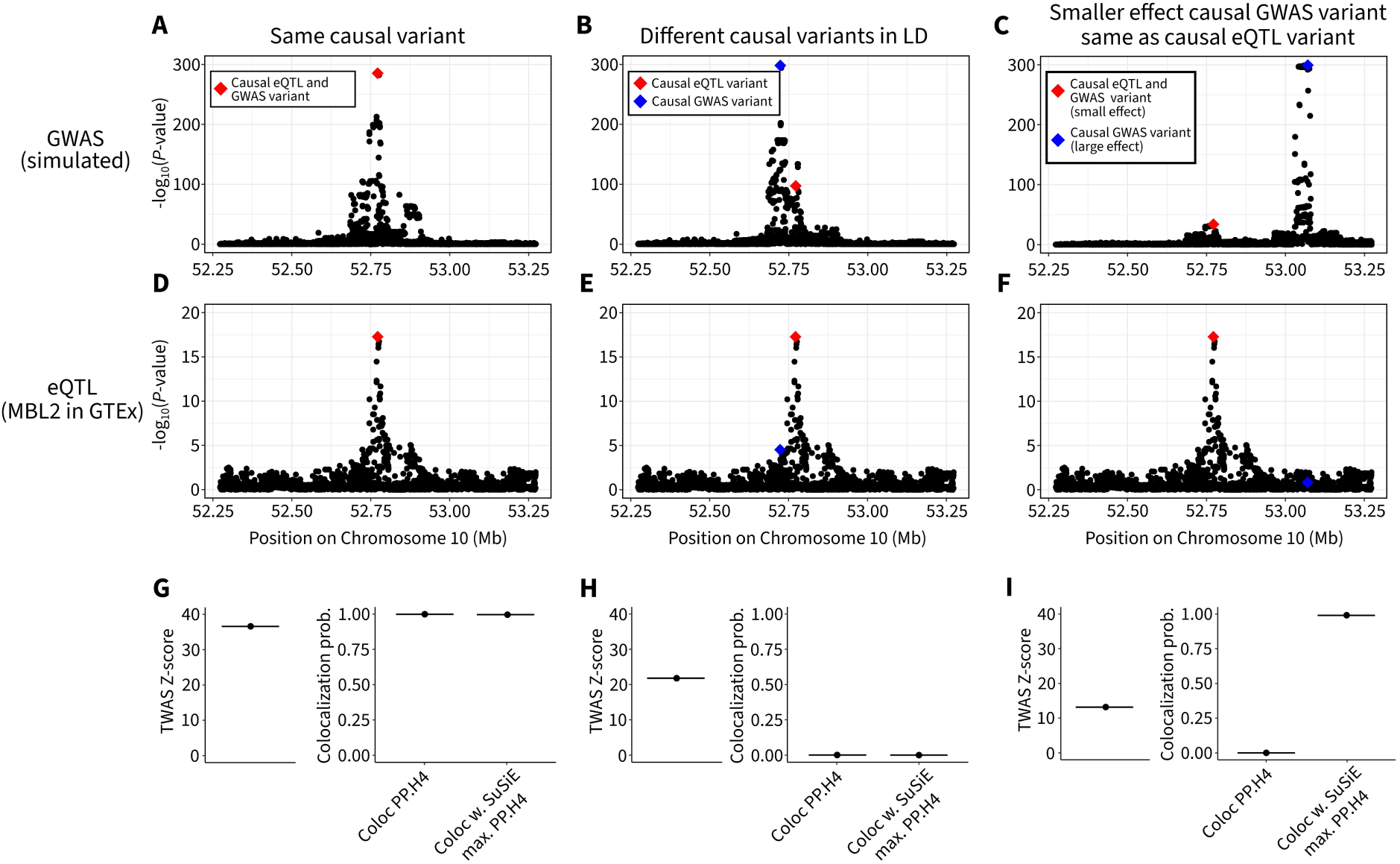
1. Simulations of TWAS and colocalization under different causal architectures. A–C, Simulated GWAS association statistics at the *MBL2* locus for three scenarios: (A) a shared causal variant between GWAS and eQTL, (B) distinct GWAS and eQTL causal variants in strong LD (*R*^2^ = 0.35), and (C) two GWAS causal variants where the smaller-effect variant is the same variant as the eQTL causal variant and the larger-effect variant is unlinked to the eQTL causal variant (*R*^2^ = 0.002). **D–F,** eQTL summary statistics for the *MBL2* locus in liver tissue. **G–I,** TWAS Z-scores and colocalization posterior probability PPH4 from coloc and coloc.susie for each scenario; for coloc.susie we show the maximum PPH4 across SuSiE credible sets. Lines in **G–I** represents the mean value across 50 simulations.

## C Appendix 3

We explored some of the discordant associations we observed between LoF burden tests and TWAS.

### C.1 *GCK* and Blood Glucose

The first example was the association between *GCK* and blood glucose levels, which had discordant TWAS signs in all tissues. *GCK* encodes glucokinase (GCK), an enzyme that catalyzes the first step in glycolysis and acts as a glucose sensor in various tissues. LoF variants in *GCK* are associated with increased blood glucose levels [5–7]. The mechanism is thought to occur via pancreatic beta cells, where the inability to detect glucose levels results in lower insulin secretion and hyperglycemia [8]. Activating mutations in *GCK* cause hypoglycemia due to increased insulin secretion [9]. Therefore, for this gene, we expect TWAS to have a negative Z-score, since increased *GCK* expression should be associated with reduced glucose levels. There is a strong GWAS signal present at the *GCK* locus, but no eQTL signal in the pancreas, resulting in no significant TWAS association in the tissue.

There were significant TWAS association in 17 tissue groups for *GCK* and glucose, all of which were in the wrong direction. Tibial nerve tissue had the most significant eQTL for this gene, so we analyzed this tissue further. In addition, we observed striking colocalization with thyroid tissue, so we analyzed this tissue as well. We identified no credible sets in pancreas, one credible set in thyroid, and one credible set in tibial nerve tissue. We concurrently identified 9 credible sets for blood glucose levels at the locus.

There was no evidence for colocalization between the pancreas Bayes factors and the GWAS signal due to a lack of eQTL signal in the pancreas (PPH2 > 0.8 for all credible sets). The credible set from the tibial nerve eQTL summary statistics had some evidence of colocalization with the second strongest GWAS signal (PPH4 = 0.51). The thyroid credible set colocalized with a high posterior probability (PPH4 = 0.99) to the primary GWAS credible set, and plotting the association summary statistics against each other shows strong evidence for the same variants underlying both signals, but in the wrong direction.

### C.2 *CXCL2* and Neutrophil Percentage

**Figure C.1.**
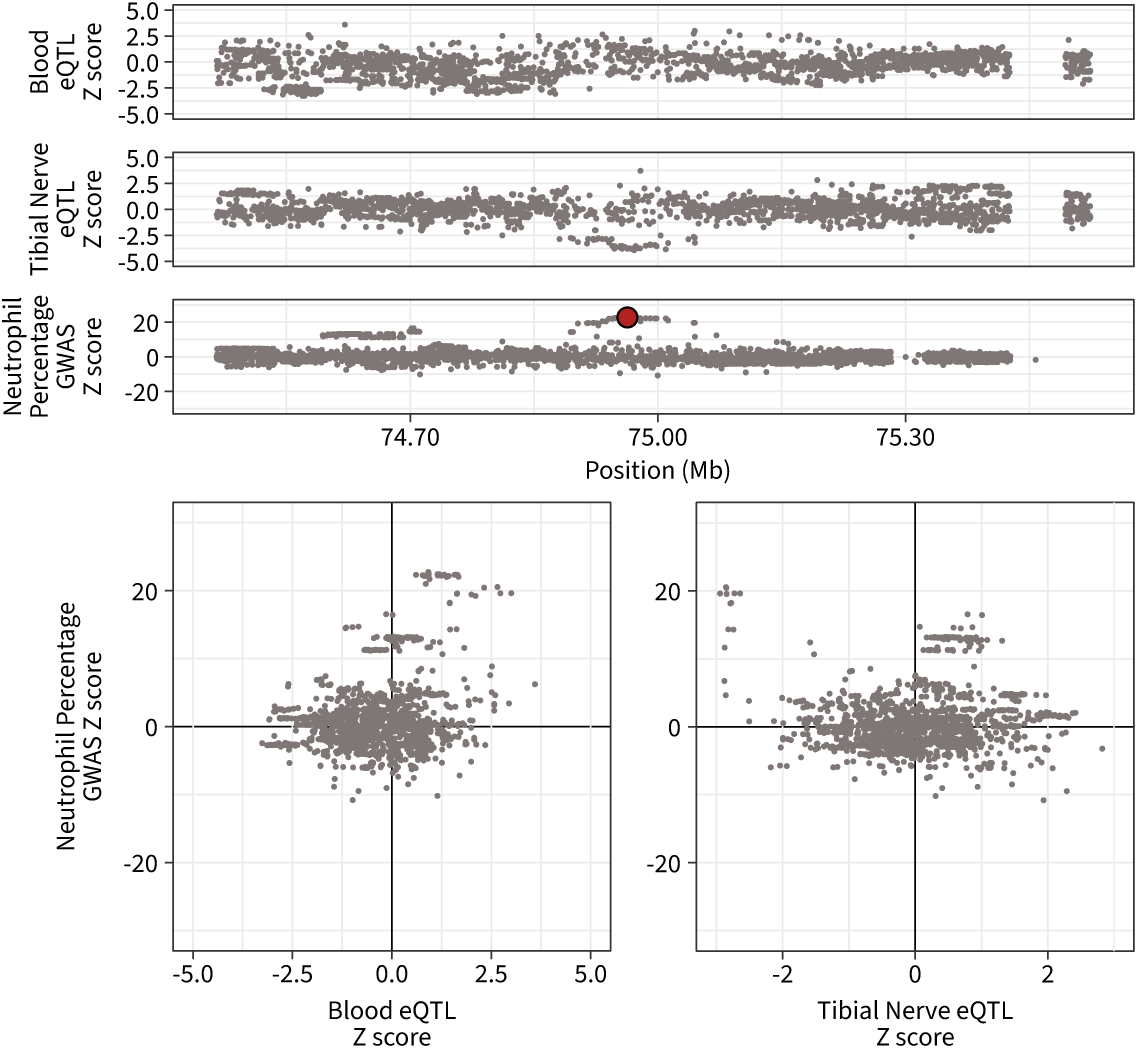
TWAS associations at the *CXCL2* Locus. *CXCL2* is a chemokine that increases neutrophil levels. In whole blood, which is the closest to a causal tissue, there is a hint of a positive relationship, which is expected. In contrast, in tibial nerve, we observe a strong negative relationship that has both colocalization signal and results in a negative TWAS Z-score.

The *CXCL2* gene encodes chemokine (C-X-C motif) ligand 2 (CXCL2), a protein that acts as a chemoattractant for leukocytes during inflammation [10]. LoF variants in *CXCL2* significantly reduce both neutrophil count and neutrophil percentage in blood [11, 12], consistent with the role of CXCL2 in recruiting neutrophils. We expect TWAS to have a positive Z-score, since decreased expression should reduce neutrophil percentage.

*CXCL2* had significant TWAS associations with neutrophil percentage in 11 tissue groups, of which only one (Small Intestine) was in the expected direction. The most significant eQTL for *CXCL2* was in tibial nerve tissue. However, processing the tibial nerve eQTL summary statistics with SuSiE failed to identify any credible sets. In contrast, SuSiE identified seven credible sets from the GWAS summary statistics for neutrophil percentage at the locus. We tested the approximate Bayes factors from the eQTL summary statistics against the credible sets from the GWAS. The primary credible set from the GWAS colocalized with the eQTL data with high posterior probability (PPH4 = 0.87). Similar to *GCK* and glucose, the summary statistics visually show strong evidence for an association, but in the wrong direction.

### C.3 *GMPR* and Mean Corpuscular Hemoglobin

**Figure C.2.**
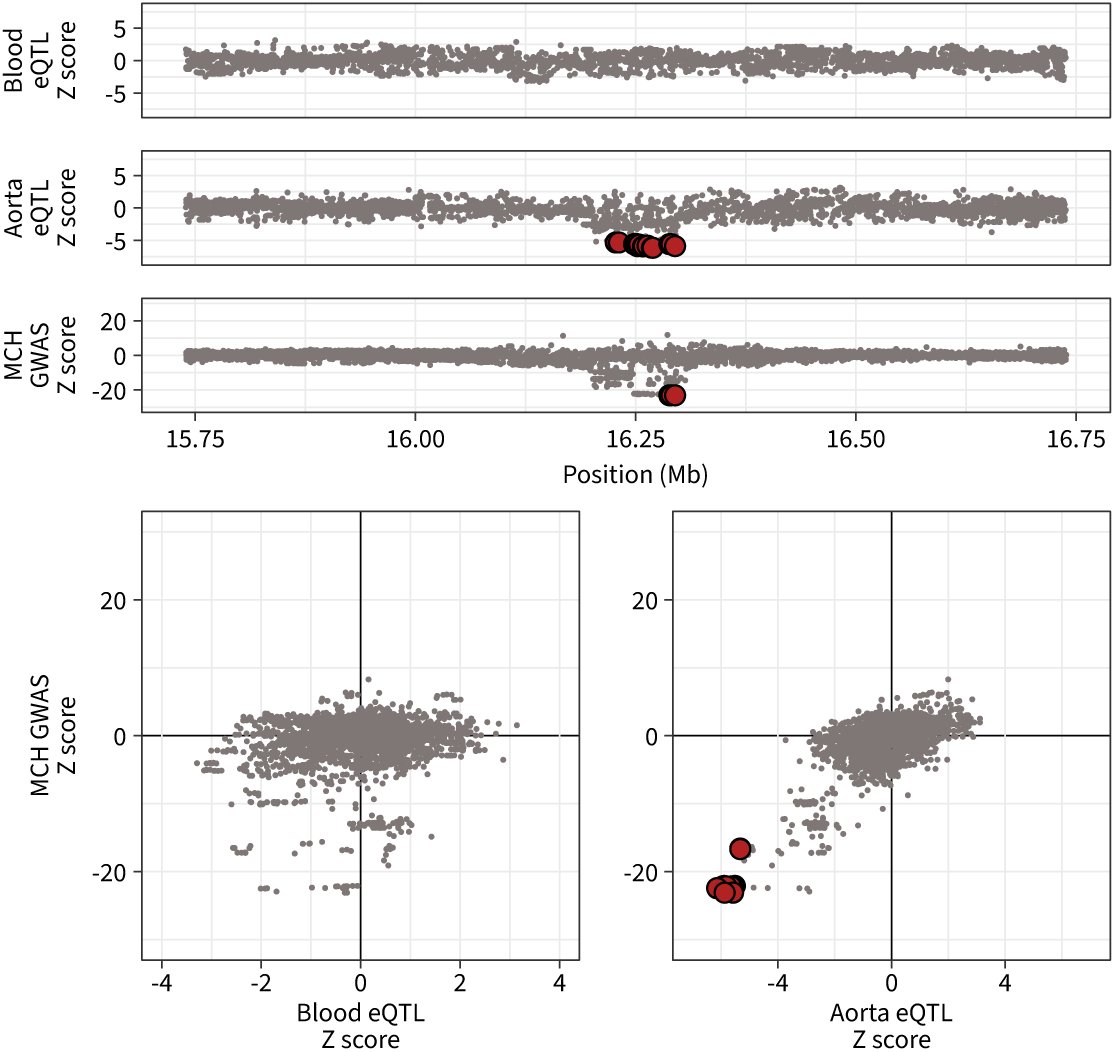
TWAS associations at the *GMPR* Locus. *GMPR* deletion is associated with increased hemoglobin concentration, so we expect TWAS associations to be negative. TWAS reports a positive Z-score in blood, but a negative Z-score in aorta. The evidence for colocalization is much stronger in the aorta.

The *GMPR* gene encodes GMP reductase (GMPR), which has previously been associated with hemoglobin levels [11–13]. LoF variants in *GMPR* are associated with increased mean corpuscular hemoglobin (MCH). GMPR may be involved in cell cycle regulation [14], which is known to have a negative effect on MCH [15]. We expect TWAS to have a negative Z-score, since decreased GMPR is associated with increased hemoglobin levels.

*GMPR* had significant TWAS associations with MCH in 14 tissue groups, of which blood, brain, and kidney showed the correct direction-of-effect. Blood is the closest tissue to the trait available in GTEx, and it is reassuring to see the correct direction-of-effect. The most significant eQTL were present in artery tissue, which were significantly associated in TWAS but in the wrong direction. We analyzed eQTL summary statistics from blood and aorta to better understand this inconsistency. We identified no credible sets in blood, and a single credible set in the aorta. We identified nine credible sets for MCH at the locus. The blood eQTL showed no evidence of colocalization, and visually did not show any evidence of association either. In contrast, the credible set from the aorta colocalized strongly with the primary GWAS credible set (PPH4 = 0.93), and the same variants from both the eQTL and GWAS summary statistics seem to explain the signal, but in the wrong direction.

### C.4 *ADAMTS17* and Height

**Figure C.3.**
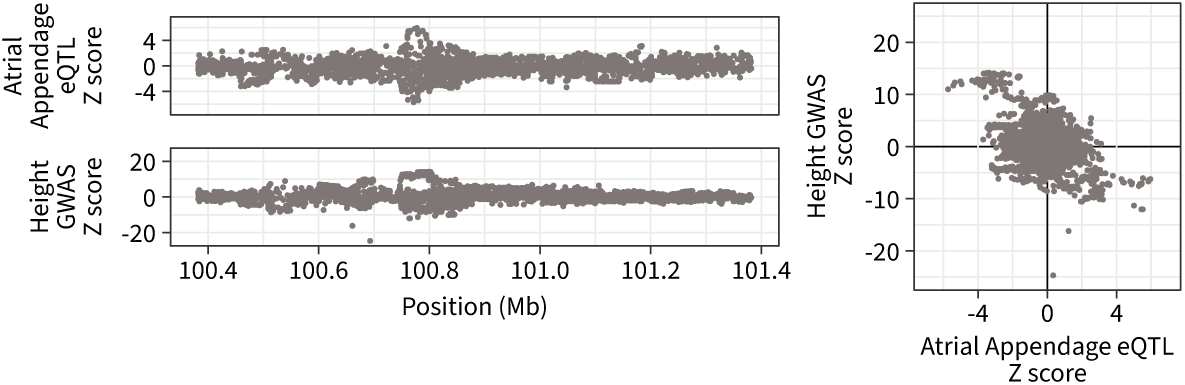
TWAS associations at the *ADAMTS17* Locus. *ADAMTS17* is a positive modulator of height, so TWAS is expected to return positive Z-scores. However, TWAS returned a negative Z-score in heart tissue, although this association would be filtered by colocalization.

The *ADAMTS17* gene encodes ADAM metallopeptidase with thrombospondin type 1 motif, 17 (ADAMTS17). LoF variants in this gene are associated with a reduction in height [11, 12]. Homozygous LoF genotypes are associated with Weill-Marchesani syndrome, which is a Mendelian disorder that presents with short stature [16]. Therefore, we expect TWAS to have a positive Z-score, since decreased expression should reduce height.

*ADAMTS17* only had significant TWAS results in two tissue groups, both of which were in the wrong direction. The strongest eQTL were present in thyroid tissue, but this model was not available for TWAS. Of the two tissue groups with models (heart and brain), the atrial appendage tissue had the most significant eQTL, which we analyzed further. SuSiE did not detect any credible sets with the eQTL summary statistics, so we proceeded with approximate Bayes factors instead. We identified ten credible sets for height at the locus, of which none colocalized with the heart eQTL summary statistics (PPH3 > 0.99 for all credible sets). This is an example of an association that would be filtered out by using colocalization or SuSiE after TWAS to filter gene-trait pairs. However, visually, the variants are strikingly correlated, which explains the TWAS Z-score.

## D Appendix 4

We used a regression strategy to test if any gene-level features explained why TWAS assigned a confidently incorrect sign at certain loci. We used four simple features: (1) the number of genes in a 20 kilobase region around the TSS, (2) the gene constraint as measured by log_10_(*s*_het_), (3) the mean recombination rate in the 1 megabase window around the TSS in the EUR superpopulation from the 1000 genomes project [17], and (4) the mean of the *cis*-heritability estimates reported in the constructed FUSION TWAS models.

TWAS performance may also be explained by the power of the GWAS analysis. For this reason, we included the ssample size of the GWAS for each trait as a covariate. Additionally, we have previously estimated trait-level monotonicity, which describes the average monotonicity of gene dosage response curves [12]. Since more non-monotone gene dosage response curves might result in more discordance of gene-level effect estimates, we also included our monotonicity estimates as covariates.

We analyzed the TWAS Z-scores that were calculated from the “filtered” gene list. We calculated the Bonferroni absolute Z-score threshold for a two-sided test based on the number of Z-scores available across all tissues and gene-trait pairs. We then used a majority vote strategy to assign a sign to each gene in its tissue group. For example, we assigned one sign to TWAS in the brain tissue group rather than using 13 brain tissues available in GTEx. This is the same data presented in Figure 5A.

We counted the total number of significant Z-scores (*N_ij_*) and the subset of those that were confidently in the wrong direction (*Y_ij_*) for the *i*th gene and *j*th trait. Then, we modeled the counts as

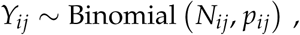

with the link function

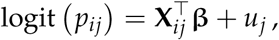

where

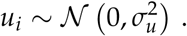

Here, **X***_ij_* represent the values of the various covariates that may explain the incorrect sign from TWAS, and **β** represents the vector of fixed effects. The random effect *u_j_* for each trait should capture any trait-specific effects on TWAS that are not explained by GWAS power or monotonicity. This is a generalized linear mixed effects model in the binomial family with a logit link function, which can be fit using the lme4 package in R. Specifically, using the formula language from lme4, we fit the model

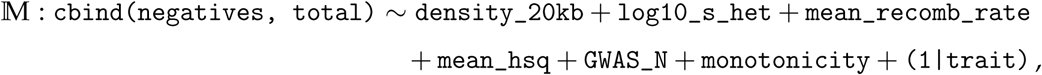

and compared it to a null model with trait-specific confounders,

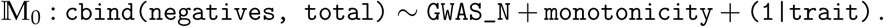

Using an analysis of variance (ANOVA), we found that M was a significantly better fit than M_0_ (*p* = 3.6 *×* 10*^−^*^4^), suggesting that gene density, gene constraint, *cis*-heritability, and mean recombination rate likely explain some of incorrect sign assignment. Using likelihood-based inference, the coefficients for gene constraint and mean recombination rate were significant in M. A unit increase in log_10_(*s*_het_) was estimated to reduce the odds of an incorrect assignment to 0.83 (*p* = 0.01), while a unit increase in log_10_(mean_recomb_rate) was estimated to increase the odds of an incorrect assignment to 1.51 (*p* = 0.001). This suggests that constrained genes tend to have fewer TWAS sign errors, and regions with higher mean recombination rate tend to have more TWAS sign errors. However, we interpreted these associations cautiously and conservatively because this analysis is based on only 383 gene-trait pairs from 186 unique genes and 68 unique traits. Therefore, we conclude that gene density, gene constraint, *cis*-heritability, and mean recombination rate may, in combination, explain some of the variation in TWAS sign inconsistency.

## Notes

### Competing Interest Statement

The authors have declared no competing interest.

